# Protein hydration and druggability

**DOI:** 10.64898/2026.07.06.736750

**Authors:** Svetlana Panasenko, Vadim Khorev, Michael Petukhov

**Affiliations:** Petersburg Nuclear Physics Institute named by B.P. Konstantinov of National Research Centre «Kurchatov Institute»

## Abstract

*A priori* assessment of target proteins’ druggability remains an unsolved problem in the field of drug development. The empirical approaches widely used to solve this problem demonstrate low efficiency. In this work, we investigated the factor of hydration of a representative set of 65 evolutionarily and structurally unrelated human enzymes in a water environment. This factor depends only on the structure of the proteins, and not on the physical and chemical properties of any potential ligands. The results show that, unlike the widely used approaches based on calculations of the accessible surface area (ASA), the content of low-entropy water molecules (LEW) in the active sites of human enzymes is systematically higher than that in other areas of their surface, including inactive cavities. Optimal criteria and a step-by-step procedure for identifying protein ligand binding sites are proposed. The proposed approach, based on the calculation of the LEW content in the first hydration layer of potentially interesting target proteins, makes it possible to evaluate their medicinal suitability even before the development of any ligands. The article also presents the results of a comparative analysis of experimental Raman spectroscopy data and the results of molecular dynamics simulations of water hydrogen bonds using three widely used water models (TIP3P, OPC3, and TIP5P) and standard algorithms for calculating hydrogen bond networks.

---

The binding of proteins to ligands is the basis of many biochemical processes, and a detailed understanding of the mechanisms of this phenomenon is an important step in the process of creating new drugs, the success rate of which, unfortunately, is still quite low. Despite significant efforts to improve the methods used, the majority of candidate compounds (∼96%) are eliminated at various stages of clinical trials^1^. Moreover, more than 60% of these cases are associated with the choice of undruggable target proteins at the earliest stages of development^2^. The situation is aggravated by the fact that almost 90% of human proteins are still considered inaccessible for therapeutic effects using low molecular weight compounds^3^. Therefore, the creation of methods for *a priori* evaluation of the medicinal suitability of target proteins *in silico* remains an unsolved challenge in the field of drug development, even with the use of modern computational methods of Structure-Based Drug Design (SBDD).

Existing computational algorithms for finding potential ligand binding sites include a wide range of approaches. Methods for identifying patterns in amino acid sequences in target proteins using machine learning models are being actively developed^3^. Structural analysis methods are also widely used, implemented in the most common software packages – ICM-Pro^4^, Schrodinger^5^ and MOE (Molecular Operating Environment)^6^, which allow detailed investigation of the geometry and physical and chemical characteristics of the protein surface. In particular, computational methods such as PocketFinder in ICM-Pro^7^, SiteMap in Schrodinger^8,9^ and Site Finder in MOE^10^ are used to identify and quantify the druggability of potential binding sites.

Generally, they identify cavities on the surface of the macromolecule and calculate an integral indicator of the druggability of the site, one of the important components of which is hydrophobicity. For example, the ICM-Pro final assessment is performed using the DLID metric based on three parameters, one of which is the fraction of non-polar solvent-accessible surface (ASA)^11^. In the SiteMap (Schrodinger), the hydrophobic contribution is calculated from the interaction energy of a hydrophobic probe with a three-dimensional Lennard-Jones potential grid^8,9^. In the Site Finder (MOE), the inner surface of the cavity is described by a set of alpha-spheres, which are then divided into hydrophobic and hydrophilic regions depending on the availability of surrounding protein atoms - donors and acceptors of hydrogen bonds^10^. Thus, despite the difference in algorithms, the hydrophobic properties of the cavity in these approaches are determined solely by the parameters of the cavity itself, while the actual behavior of water molecules forming a dynamic network of hydrogen bonds remains out of the scope of consideration.

It is known that during protein-ligand binding, the hydrophobic effect is associated with the displacement of low-entropy water molecules in direct contact with nonpolar areas of the protein surface from the binding sites into bulk water (hereinafter such water molecules will be denoted as low-entropy water molecules or LEW). In bulk water at normal physiological temperatures (20-37°C), each water molecule forms from 3 to 4 hydrogen bonds with neighboring water molecules, forming mobile networks of intermolecular hydrogen bonds^12^. In the cavities of the active sites of globular water-soluble proteins, the ability of water molecules to form hydrogen bonds is significantly reduced. Thus, the quantitative criterion for LEW in this work is the formation of 1, 2 or no hydrogen bonds. The nonpolar groups of the protein surface are unable to form hydrogen bonds, and steric constraints prevent optimal orientation of water molecules even relative to those polar groups located in the active centers. As a result, water molecules in the active centers of proteins may have fewer than three hydrogen bonds and, consequently, lower entropy. It is of interest that the displacement of each LEW from a protein active site may produce a gain in free energy of up to 2.0 kcal/mol^13^. Fig. 1 shows a typical example of the distribution of LEW on the surface of the enzyme human phytanoyl-CoA dioxygenase. As seen in this figure, the content of LEW in the zone of the active center of this enzyme is significantly higher than that in other areas of the protein surface.

**Fig. 1.**
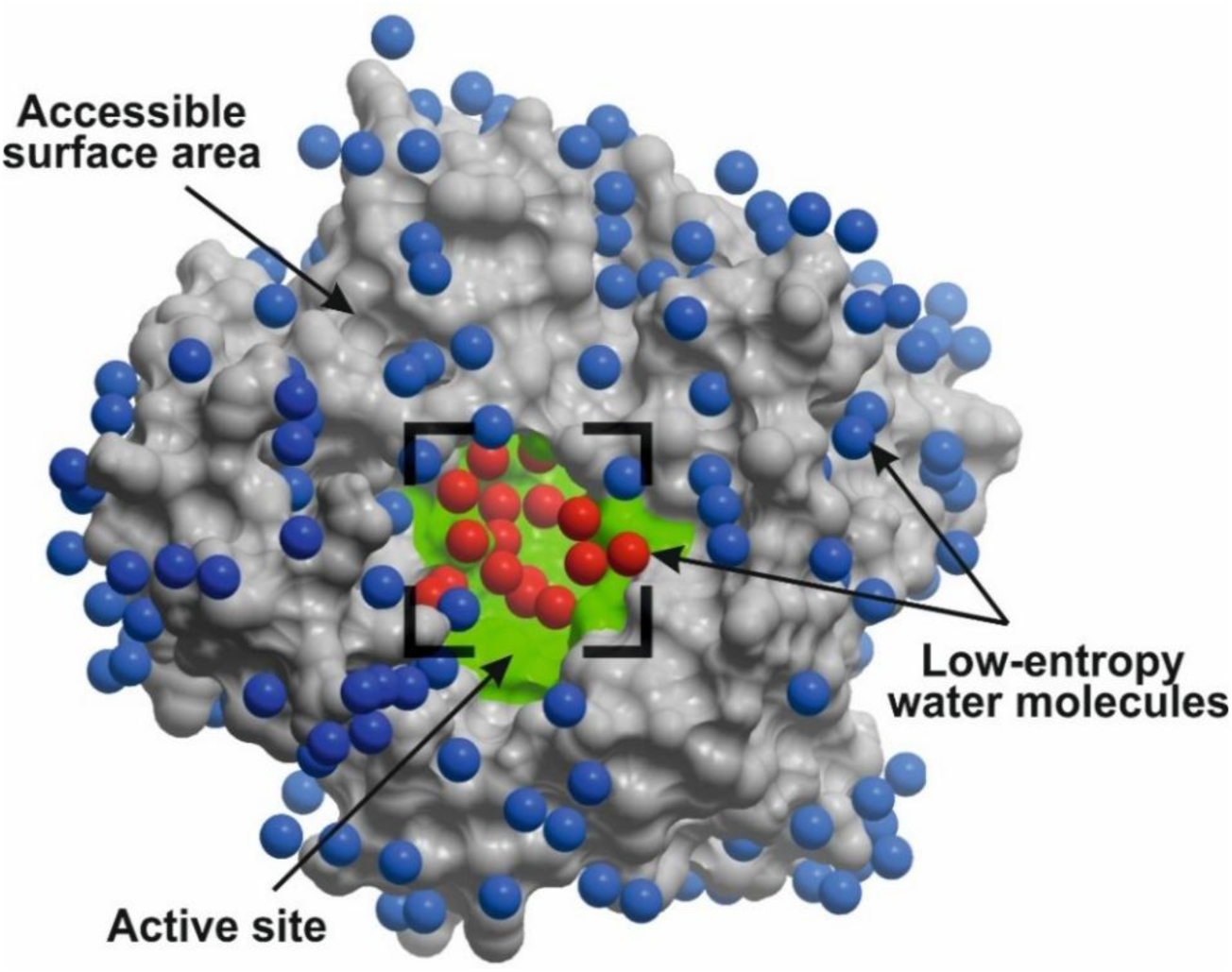
A frame from the MD trajectory of human phytanoyl-CoA-dioxygenase (PDB ID: 3OBZ), showing a typical LEW distribution (TIP5P) on the protein surface. The surface of the active site is shown in green. The red spheres indicate the LEWs in the region of the active site; the blue spheres indicate the LEWs on the rest of the protein surface.

It has been shown that the level of hydrophobicity, estimated as the average value of the hydrophobicity index of amino acid residues forming a cavity, does not allow for distinguishing between active centers and inactive cavities^2^. Also, there is a series of ASA-based continuous hydration models of proteins^11^ that are widely used for assessing the level of hydrophobicity of the protein surface, including active sites. These models are derived from experimental data on the equilibrium thermodynamics of the transfer of small organic compounds from vapor or cyclohexane to water^14^, assuming a simple proportionality between the free energy of hydration and atomic ASA. However, in the case of globular proteins, they often produce contradictory results^15^. Moreover, it has been shown that protein hydration differs significantly from the hydration of small organic compounds both in terms of typical atomic ASA values and atomic hydration parameters (ASP), which themselves are non-linearly dependent on ASA^16^. In addition, the polar/charged and hydrophobic groups in the protein active sites are closely adjacent in space, and their hydrate shells often overlap. Therefore, the ability of neighboring hydrophobic sites to bind LEW inevitably depends not only on the total area of the hydrophobic surface of the cavity, but also on the relative location of neighboring polar/charged and nonpolar groups. Consequently, the contribution of each unit of ASA to the binding of LEW is not constant and it is impossible to simply sum up the areas of several small hydrophobic sites, expecting a proportional amount of LEW bound on this surface.

Other protein characteristics, such as the interaction energy of the hydrophobic probe and the classification of alpha spheres, which are currently used to assess the hydrophobicity of certain protein regions, also do not take into account the cooperative nature of LEW binding in potentially interesting protein binding sites.

Modern methods of molecular dynamics (MD) of proteins in an aqueous environment are capable of obtaining statistics on the distribution of LEW on any part of the protein surface. In this paper, we investigated the content of LEW (water molecules with 0-2 hydrogen bonds) in active centers, inactive pockets, and the rest of the surface of a representative set of 65 human enzymes from the PDB. Since this requires precise knowledge of the number of hydrogen bonds individually for each water molecule in the first layer of protein hydration, we analyzed the abilities of various water models and algorithms for calculating hydrogen bonds to reproduce the available experimental data on the structure of liquid water, which were obtained using Raman spectroscopy^12^.

## Identification of hydrogen bonds

Currently, there are several different molecular models of water molecules, as well as computational algorithms for identifying hydrogen bonds in the molecular modeling of proteins and their complexes with water and other ligands. In this work, we analyzed 3 water models (TIP3P, OPC3 and TIP5P) and 4 standard algorithms for calculating hydrogen bond networks used in widely used software packages (AMBER^17^, GROMACS^18^ and ICM-Pro^4^).

The results of this comparative analysis shown in Fig. 2 demonstrate significant differences in the ability of all tested approaches to reproduce the dependence of the fraction of water molecules having four hydrogen bonds in bulk water over the temperature range 0-100°C^12^.

**Fig. 2.**
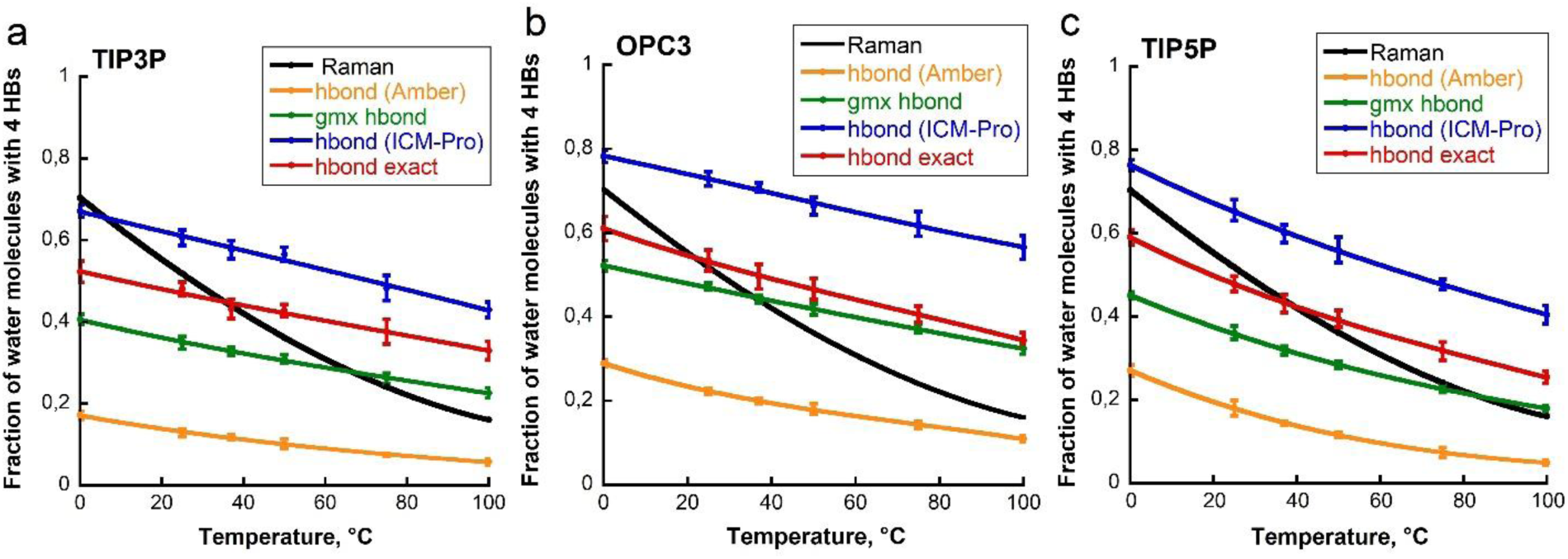
Temperature dependence of the fraction of water molecules in bulk water with 4 hydrogen bonds: comparison of water molecule models and hydrogen bond identification algorithms. Panels **a**, **b** and **c** show results for TIP3P, OPC3 and TIP5P respectively. The experimental data of Raman spectroscopy^12^ are shown in black. The colored lines correspond to the results of MD simulations using the following algorithms of HB identification: ‘hbond’ of the AMBER software package (shown in amber), ‘gmx hbond’ of the GROMACS package (shown in green), ICM-Pro’s ‘hbond’ (shown in blue) and ‘hbond exact’. The error bars correspond to the standard deviations obtained from 10 ns MD simulations of a water box.

As expected, the best combination of the water model/HB identification method turned out to be the TIP5P/’hbond exact’ (ICM-Pro) pair, which has the most detailed characterization for both the structure of water and the method of hydrogen bond identification. In contrast to the simplified three-point TIP3P and OPC3 water models (Fig. 2, panels a and b), the five-point TIP5P model (Fig. 2, panel c) demonstrates a nonlinear temperature dependence close to the experimentally observed curve of temperature dependence, especially in the biologically interesting temperature range of 20-37°C (RMSE = 0.077).

## The LEW content is a key indicator of druggability of potential target proteins

In this work, we analyzed the LEW content in three types of surfaces of a representative set of 65 evolutionarily and structurally unrelated human enzymes, a list of which is presented in Table 1 of the Methods section. Calculations were performed for: 1) active sites, 2) structurally similar inactive cavities, and 3) the rest of the protein ASA. The hydration analysis of each of the 65 proteins without ligands in a periodic water container was performed based on MD trajectories (10 ns). 10 frames with MD time steps of 1 ns were obtained from each MD trajectory, which were subsequently used to calculate the content of LEW and hydrophilic/hydrophobic ASA and to perform statistical analysis.

**Table 1.** A representative set of 65 structurally and evolutionarily unrelated human proteins from the PDB.

| EC number | Quantity | PDB ID |
| --- | --- | --- |
| **Oxidoreductases (1.x.x.x)** | 18 | 2GCG, 2I99, 2PLA, 2YDE, 3F3S, 3OBZ, 3RGK, 4BKP, 6N0K, 6USN, 7VJV, 8ABX, 8ELL, 8GRZ, 8VHD, 8WGN, 9D71, 9EZT |
| **Transferases (2.x.x.x)** | 26 | 1W7N, 1XW6, 2AOX, 2CH6, 2JEV, 2PQT, 2XX3, 3JUC, 3RIC, 4C2K, 4KN6, 5JZV, 6HGS, 6P7Z, 6PNO, 6RV0, 6T8Q, 6YAW, 7JR8*, 7ZSP, 8E8P*, 8PYW, 8UG3, 8WR2, 9FDN, 9HJC |
| **Hydrolases (3.x.x.x)** | 17 | 1ONI, 1ZS9, 2HW4, 2HZP, 2WEF, 3VKK, 4NC5, 5L0A, 5QIZ, 5VXA, 6FIT, 6IUX, 6Q6J, 7B64, 7UW6, 8EFG, 8TK5 |
| **Lyases (4.x.x.x)** | 1 | 8ORA |
| **Isomerases (5.x.x.x)** | 3 | 7JR8*, 8E8P*, 9FFC |
| **Ligases (6.x.x.x)** | 2 | 3HY3, 5F6Y |

\* 7JR8 and 8E8P have a dual classification (Transferase/Isomerase)

Fig. 3a shows the differences in the average content of LEW having < 3 hydrogen bonds for the three types of protein surfaces. The average proportion of LEW was 0.46 ± 0.06 in active centers, 0.42 ± 0.04 in inactive cavities, and 0.38 ± 0.02 on the rest of the protein surface. In approximately 68% of all studied proteins, the proportion of LEW in the active sites is higher than in their inactive cavities and on the surface not included in the first two types. Estimates of the statistical significance of the observed differences in the average values of LEW using the nonparametric Mann-Whitney U-test criterion showed that the proportion of LEW in active centers is statistically significantly higher than on the rest of the protein surface, including inactive cavities (p < 0.0001). The difference of LEW content between the inactive cavities and the exposed surface is also statistically significant (p < 0.0001).

**Fig. 3a.**
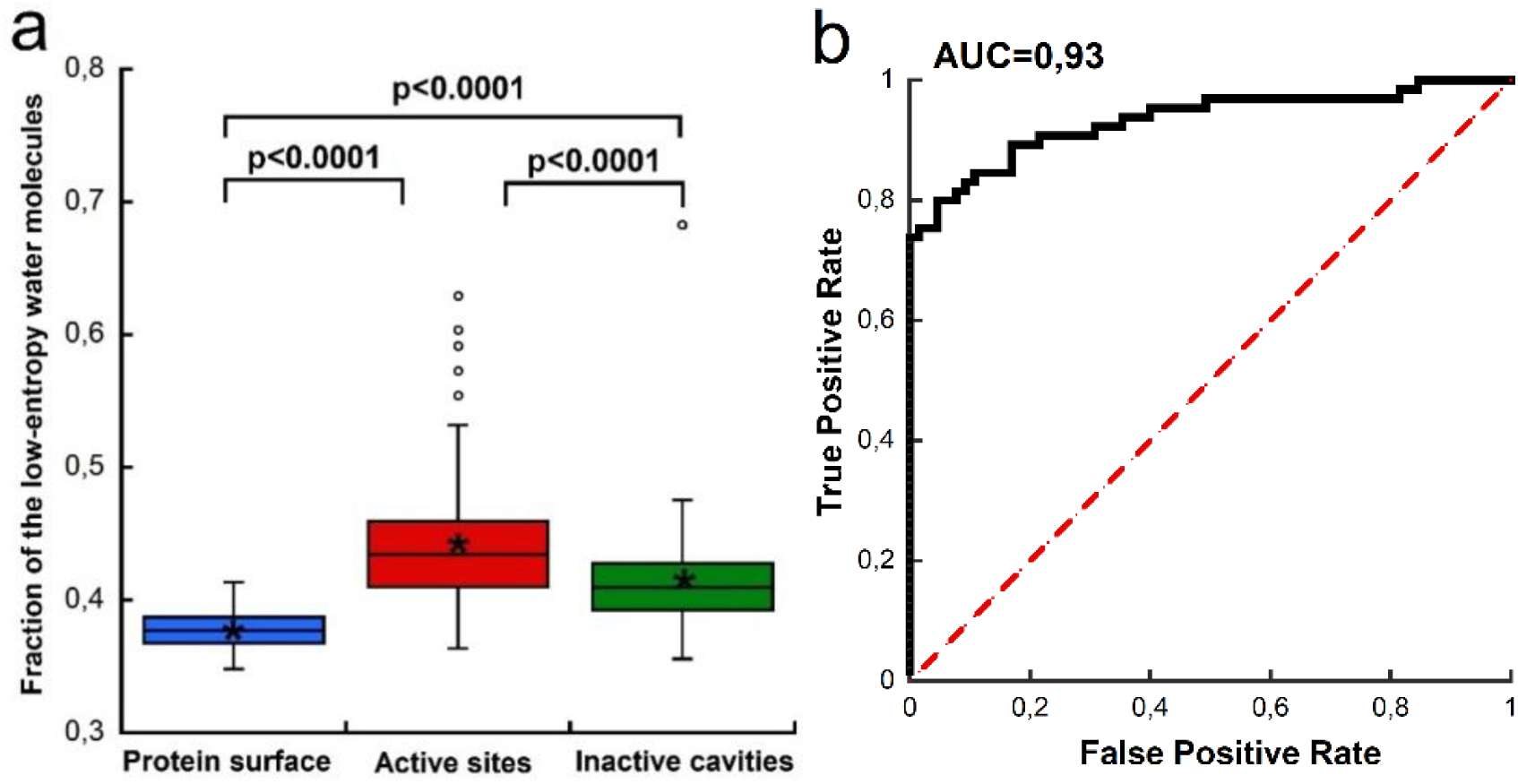
Distribution of the low-entropy water (LEW) fraction in different functional zones of the surface of the 65 proteins of the representative set. The “*” signs and horizontal lines inside the boxes indicate the average and median values, respectively. The color boxes contain data from the first to the third quartile (∼50%). The vertical lines show the boundaries of the minimum and maximum values within 1.5 quartile intervals from the boundaries of the corresponding boxes. The dots show anomalous values of the LEW content that fall outside the distributions; **b,** ROC analysis of statistical significance for optimal separation of active sites and exposed protein surfaces based on the LEW content in their first hydration layer. Calculations were performed using a cutoff value of 0.405 for the LEW content in protein active sites, which provides the maximum Youden J-index^19^ for the representative set of 65 human proteins.

It is of interest, most of the anomalous deviations shown in Fig. 3a are observed for active sites and are always directed towards an increase in the LEW content, which is a clear indication of the important role played by active site hydration in the functional activity of these enzymes. The solubility of proteins in water is a very important parameter of their biological activity. Too high a content of LEW on a protein surface can lead to undesirable aggregation. This probably explains the lower values of the LEW content in inactive cavities and in the rest of the protein surface, as well as the small spread of their hydrophobicity indicated by the colored boxes shown in Fig. 3.

## Identification of active sites of human proteins

In terms of the practical application of the obtained results for identifying potentially interesting ligand-binding sites, not only the optimal cutoff level for the LEW content but also an assessment of the expected level of statistical significance of this approach are crucial. To select the optimal cutoff level, we used a standard approach based on calculating the maximum value of the Youden J-index^19^ for two protein surface types from a representative set: known active sites of proteins containing ligands and the remaining protein surface not included in any empty cavities. This choice is explained by the fact that it is impossible to be completely sure that empty cavities, often found in the spatial structures of proteins, are indeed unable to bind ligands that may have been present in vivo but were absent in crystallographic experiments. This eliminates false-negative identifications of active sites and thus significantly increases the reliability of the initial data as a whole. With regard to false-positive identifications of protein active sites, choosing a cut-off level for the LEW content corresponding to the maximum of the Youden J-index maximizes the number of correct identifications and minimizes the number of false-positive ones.

To quantitatively assess the expected level of statistical significance of the approach for identification of protein active sites, a ROC analysis was performed. The ROC curve presented in Fig. 3b indicates a very high ability of the LEW content in the first hydration layer of proteins to identify their active sites (AUC = 0.93), while the optimal cutoff level for the LEW content was 0.405 with a separation sensitivity of 0.800 and a specificity of 0.954.

## LEW content and ASA-based protein hydrophobicity

Traditionally, the level of hydrophobicity of the protein surface is estimated by the content of hydrophobic ASA, assuming a simple proportionality between the free energy of hydrophobic interactions and protein ASA during ligand binding. Although this proportionality has been proven experimentally for small organic compounds, this does not appear to be the case for globular proteins^16^. However, an interesting question remains as to whether ASA-based estimates of protein surface hydrophobicity can serve as an indicator of potential binding sites for any ligands?

Fig. 4a shows the results of these calculations for the same three types of surface areas of the 65 human enzymes. The average proportion of the hydrophobic surface was 0.61 ± 0.09 in active centers, 0.59 ± 0.07 in inactive cavities, and 0.57 ± 0.03 on the exposed protein surface, ∼1.5 times higher than that of the LEW calculations shown in Fig. 3a. Although the active sites of proteins are on average more hydrophobic than the rest of the protein surface (p = 0.0065), the difference in hydrophobicity between the active sites and inactive cavities based on ASA is statistically insignificant (p = 0.2452). Thus, an important role in binding protein ligands is played not by the total area of the hydrophobic surface, but by the spatial structure of the active sites, which limits the number of hydrogen bonds between water and protein compared to bulk solvent and the cooperativity of LEW binding. This also suggests that there is no correlation between the hydrophobicity of the ASA-based surface and the content of LEW in the first hydration layer of proteins. The data in Fig. 4b and Supplementary Fig. 1 confirm this assumption. Similar results were obtained for inactive cavities and the rest of the protein surface.

**Fig. 4a.**
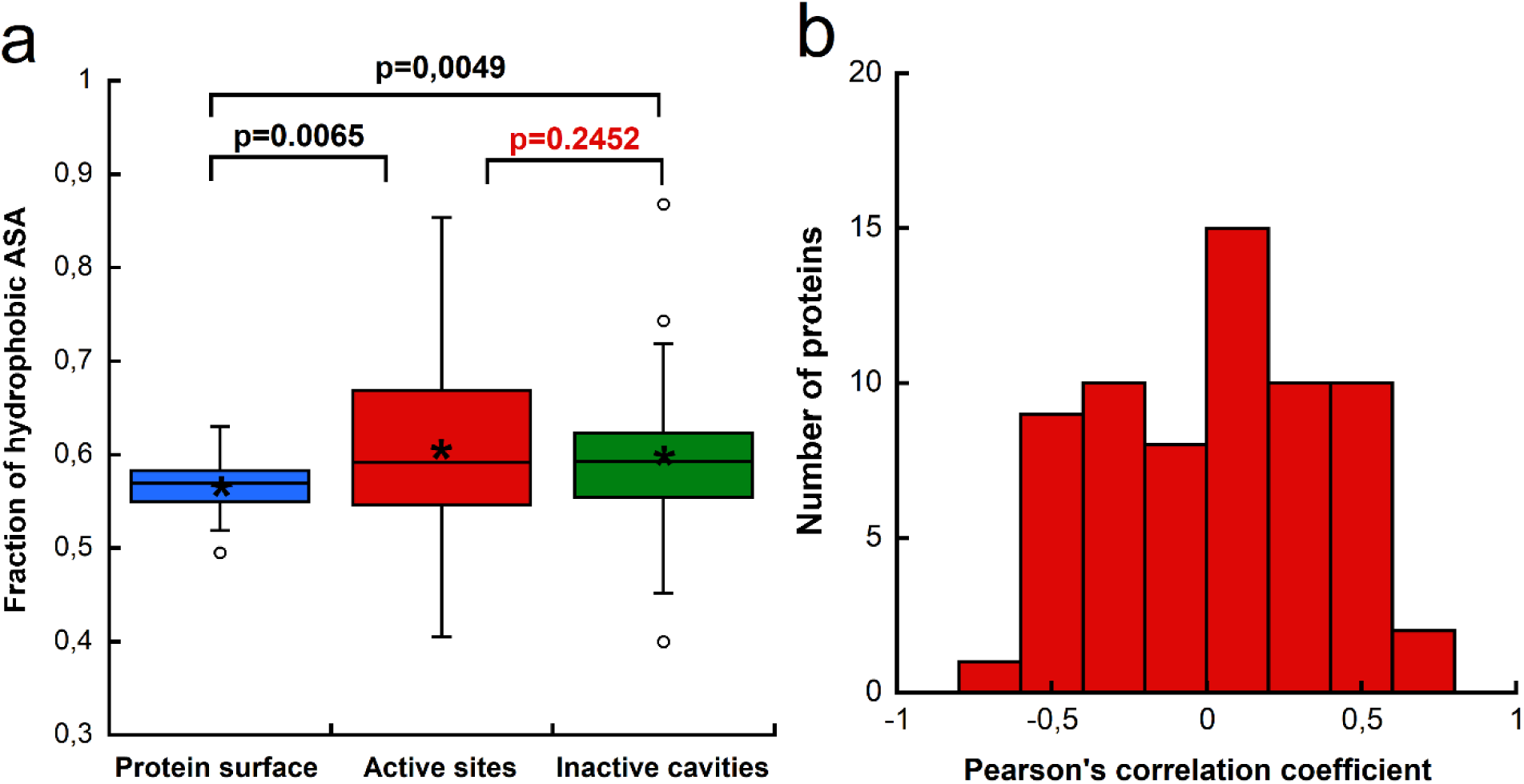
Distribution of the fraction of hydrophobic solvent-accessible surface (ASA) in active sites, inactive cavities, and on the exposed surface of the 65 proteins of the representative set. Notation is the same as in Fig. 3a**; b,** Distribution of Pearson correlation coefficients between the fraction of low-entropy water molecules (LEW) and that of hydrophobic ASA for the active sites of 65 proteins. For each protein, the correlation coefficient was calculated using 10 evenly selected frames of the molecular dynamic trajectory. The mean ± standard deviation was 0.03 ± 0.34. The distribution is almost symmetrical with respect to zero. A similar pattern of distributions, confirming the absence of correlation, is also observed for inactive cavities and the rest of the exposed surface (Supplementary Fig. 1).

## Discussion

The functional activity of enzymes and other proteins requires their ability to bind ligands in water. To achieve this, a key requirement is an increased content of LEW at their binding sites. Initially, evolutionary pressure seems to have affected the structure of not only the final active center, but also other externally accessible pockets having the necessary dimensions for ligand binding. Furthermore, it cannot be ruled out that at least some of the inactive empty cavities present on the surface of many proteins may bind ligands and play an important role in the biological activity of these proteins *in vivo*. This probably explains the intermediate position of inactive pockets on the LEW content scale between the active protein sites and the rest of the protein surface.

Thus, the analysis of the LEW content in protein hydration opens the way to a more accurate quantitative assessment of the contribution of the hydrophobic effect to the free energy of ligand binding. Unlike this approach, the active site amino acid compositions or formal hydrophobicity based on ASA do not allow for a statistically reliable separation of functionally active protein sites from inactive cavities. The LEW content is directly related to the structural and dynamic properties of proteins and does not depend on the structure and physico-chemical properties of potential ligands. The increased content of LEW in the protein active sites is a necessary, but not a sufficient, condition for binding specific ligands. Other factors such as ligand size, shape, charge, composition and location of HB donors and acceptors also play an important role. However, the approach proposed in this work can be used as an indicator for assessing the ability of potentially interesting target proteins to bind ligands even before the start of their development. Specifically, once a set of cavities has been identified on the surface of a potential target protein, applying the threshold LEW fraction of > 0.405 makes it possible to determine which of these pockets are capable of binding small-molecule compounds, and which are entirely unpromising in this regard.

## The step-by-step algorithm for identification ligand binding sites of proteins

**1. Identification of protein cavities.** Search for all available cavities on the surface of a protein under study using standard geometric approaches such as icmPocketFinder, SiteMap, Site Finder or similar.
**2. Modeling of the molecular dynamics** of an apo-form protein in a TIP5P water environment.
**3. Identification of the first layer of protein hydration.** The first layer of protein hydration includes water molecules, in which the distance from the oxygen atom to the nearest heavy protein atom does not exceed 3.4 Å for nitrogen and oxygen atoms; 3.7 Å for carbon atoms and 3.8 Å for sulfur atoms.
**4. Calculation of the number of hydrogen bonds of water molecules.** The number of hydrogen bonds for all water molecules of the first hydration layer is carried out using the “hbond exact” algorithm^20^ at an energy cutoff threshold of 1.8. Low-entropy (LEW) molecules are considered to have 0, 1 or 2 hydrogen bonds with a protein or other water molecules of a periodic water box. The LEW content is calculated as the ratio of the number of low-entropy molecules to the total number of water molecules in the first hydration layer of a potentially interesting area of the protein surface. The obtained values of the LEW content are averaged over all selected MD frames.
**5. Applying the optimal cutoff threshold.** Cavities for which the average LEW content exceeds the minimum cutoff threshold of 0.405 are considered as potentially capable of binding low molecular weight organic compounds. Moreover, the higher the LEW content, the higher the ability of the corresponding protein site to bind ligands suitable in size, structure, and other physico-chemical properties.

## Methods

### Comparative analysis of water models and hydrogen bond identification algorithms

MD modeling (NPT) was carried out for a truncated octahedral water box with a minimum distance from the central water molecule to the boundary of the box of 20 Å. MD time for each run was 10 ns for each temperature in the range from 0 to 100°C with increments of 10°C plus 37°C (human body temperature), which corresponds to the available data of Raman spectroscopy^12^.

Since there are a number of different water models and methods for identifying intermolecular hydrogen bonds of proteins in an aqueous environment, choosing the most appropriate methods for analyzing protein hydration is not an easy task. Therefore, we used MD simulations of water boxes with 3 different water models (TIP3P, OPC3 and TIP5P) as implemented in the Amber-20 software package^17^. Unlike TIP3P and OPC3, the TIP5P model includes two additional interaction centers corresponding to lone pairs of electrons in the molecular structure of a water molecule. This allows for a more accurate reproduction of the spatial structure of intermolecular contacts of water molecules.

Since the results of the analysis of hydrogen bond networks depend not only on the solvent model, but also on the method for determining hydrogen bonds, we also used 4 widely used algorithms for hydrogen bond identification, namely: ‘hbond’ (AmberTools)^21^, ‘gmx hbond’ (GROMACS)^22^, ‘hbond’^4^ and ‘hbond exact’^20^ (both are from ICM-Pro software package).

In ‘hbond’ from the AmberTools package, geometric criteria are applied, and the hydrogen bond is determined while meeting the conditions for the distance between the donor and the acceptor of < 3.0 Å and the acceptor-hydrogen-donor angle ≥ 135°. The ‘gmx hbond’ (GROMACS) uses the same approach, however, the distances and angles are slightly different – < 3.5 Å and ≥ 135°, respectively. The ICM-Pro software package^4^ uses two algorithms for calculating hydrogen bonds – ‘hbond” uses a simplified criterion based only on the distance between the proton and the acceptor < 2.5 Å; ‘hbond exact’ uses a more complex approach based on the calculation of the interaction energy between the hydrogen atom and the lone electron pair of the acceptor^20^. The interaction energy is estimated taking into account the location of the centers of the lone electron pairs, and a continuous function combining angular dependence with exponential attenuation as the atoms move away from each other. This allows for a comprehensive evaluation of hydrogen bonds by calculating a single value of the interaction energy that depends simultaneously on distance and angle. In this paper, the ‘hbond exact’ algorithm was used in our calculations with a calibration parameter of the energy of the hydrogen bond cut-off threshold of 1.8, which ensures the best match between the calculation results and experimental data.

### A representative set of 65 evolutionarily and structurally unrelated human enzymes

A representative set of evolutionarily and structurally unrelated human enzymes was selected from the PDB database^23^. The initial search covered the experimental structures of human enzymes of 6 main classes, presented both in free form and as part of complexes.

Representatives of the 7th class (translocases) were not considered, since they are membrane proteins, whereas our analysis included only water–soluble proteins localized in cell compartments with a neutral pH of ∼7.0-7.4^24^. The spatial structure of the proteins was determined by X-ray crystallography with a resolution of < 3.0 Å. Complexes containing nucleic acids, as well as proteins with missing fragments of a polypeptide chain with a length of more than five amino acids in a row were excluded. The size of the multimeric complexes was limited to six polypeptide chains. The presence of an open active site was also a mandatory criterion.

This ensured that the selected targets were in a conformation ready for binding without large-scale structural rearrangements. At the final stage, proteins with amino acid sequence identity >25% were removed from the representative^25^.

To restore the structural integrity of the proteins, their sequences from the PDB were aligned with their complete sequences taken from the UniProt database^26^. Based on the alignment of protein sequences, the missing fragments of their structure were restored using the standard protein regularization protocol of the ICM-Pro software package^4^. Table 1 shows a list of 65 structurally and evolutionarily unrelated human proteins from the PDB, obtained as a result of the above-described multi-stage selection.

To assess the representativeness of the selected set of 65 human enzymes, we compared their EC number distribution with that of known human enzymes in the UniProt database (18,374 entries, version as of April 2026). Fig. 5 shows the results of the correlation analysis. The Pearson correlation coefficient between the two datasets was R=0.90, indicating a high similarity between the two distributions. The most significant difference relates to the class of oxidoreductases, the proportion of which in our data set turned out to be about twice as high as in UniProt. It is noteworthy that during the selection of enzymes, we did not make any arbitrary adjustments in the composition of the enzymes in order to formally improve compliance with the reference distribution. Thus, at the qualitative level, the representativeness of the protein set is fully determined by the presence in the set of selected 65 proteins of all classes of water-soluble human enzymes from the PDB in proportions close to those that characterize the general sample of annotated human enzymes presented in the more complete UniProt database.

**Fig. 5.**
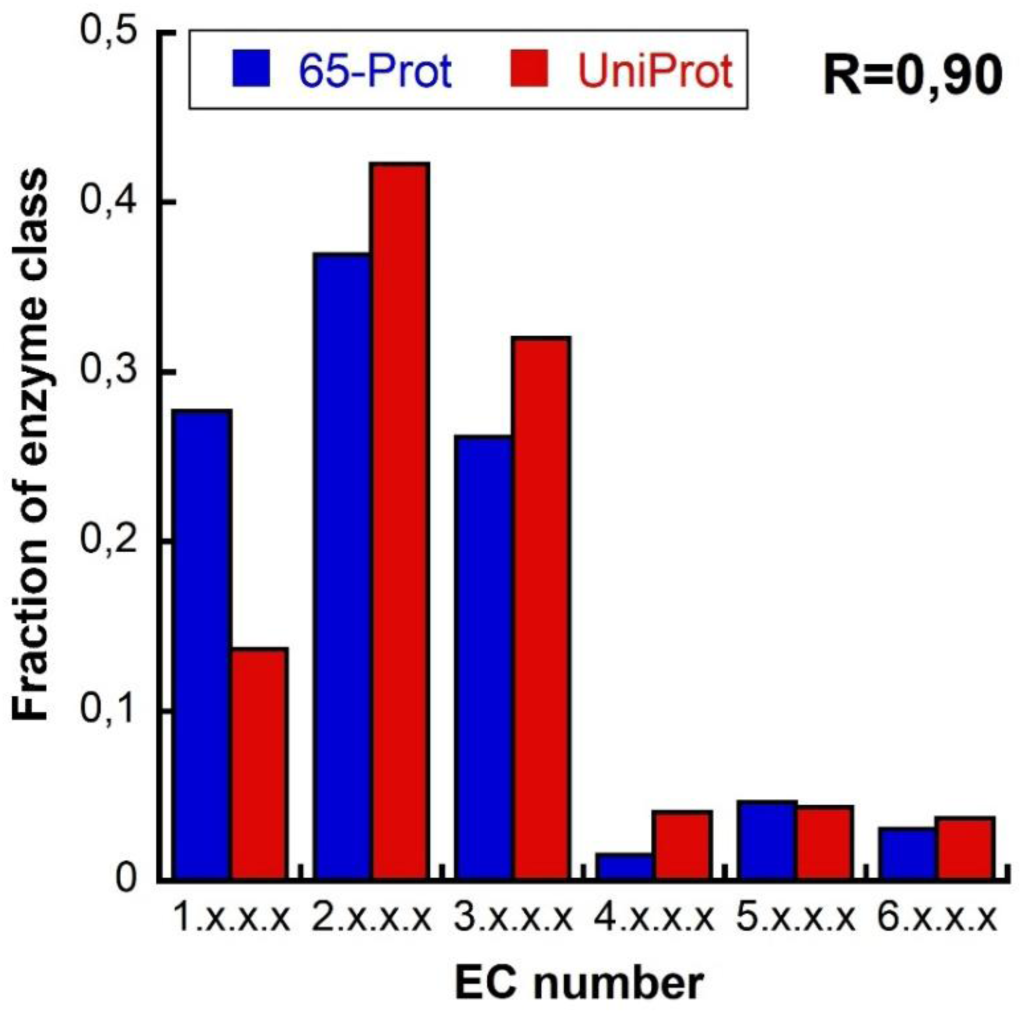
The distribution of enzyme classes in the representative set of 65 human enzymes and that of all human enzymes present in the UniProt database.

### MD simulations

MD simulations of water and the 65 selected proteins in a periodic water box was carried out with a duration of 10 ns using the Amber-20 software package and the ff14SB force field.

For proteins, MD simulations were carried out in a truncated octahedral periodic water box filled with TIP5P water molecules with a minimum distance of 12 Å from the protein atoms to the cell boundaries. The ionic composition was prepared following the SPLIT rule^27^: first, the required amount of counterions (Na⁺ or Cl⁻) was added to fully compensate for the total charge of the protein, after which the same amount of Na⁺ and Cl⁻ ions was added so that the volume concentration of NaCl was 0.15 M. Thus, the total charge of the system was zero, and the ionic strength corresponded to the physiological conditions. The MD trajectories were obtained at a physiological temperature of 310 K (37°C). For the statistical analysis of hydrogen bonds, 10 equidistant frames were obtained from each trajectory with a time step of 1 ns, which were subsequently used to identify low-entropy water molecules (LEW), as well as to calculate protein ASA and the ASA-based protein hydrophobicity.

### Identification of low-entropy water molecules

To identify the LEW on the surface of the studied proteins, we used a software script implemented in the ICM-Pro shell language^4^. The analysis was performed for 10 equidistant frames of each MD trajectory. The surface of each protein was divided into three functional zones: the active center, inactive cavities, and the rest of the exposed surface. The icmPocketFinder utility^7^ was used to divide the surface into analyzed areas. The algorithm identifies potential binding regions based on the calculation of the Lennard-Jones potential on a spatial grid. The default threshold value threshold = 4.6 was used to delineate the boundaries of the cavities. As a result, a set of cavities was obtained for each MD frame. One of the cavities was classified as an active site based on the position of the ligand in the protein structure or in case of its absence, according to the binding residues annotated in the UniProt database. The other identified cavities were considered inactive. The protein surface outside the boundaries of the cavities was assigned to the rest of the exposed surface.

The molecules of the first hydration layer were isolated based on the threshold distance <3.4 Å between the oxygen atom of the water molecule and a nearest heavy protein atom. The choice of this distance was based on the experimental data on the size of the first hydration layer of proteins^28^. A distance of 3.4 Å was used for nitrogen and oxygen atoms, covering the region of formation of hydrogen bonds. A threshold distance of 3.7 Å was used for protein carbon atoms, which is a typical value for van der Waals interactions at protein nonpolar areas. The threshold distance for sulfur atoms was 3.8 Å, due to 0.1 Å difference between the Van der Waals radii of sulfur and carbon.

The number of hydrogen bonds was calculated using the ‘hbond exact’ algorithm as implemented in the ICM-Pro software package for all water molecules of the first hydration layer of all proteins of the representative set. According to the definition adopted in this work, water molecules with 0, 1 or 2 hydrogen bonds were considered as LEW. It is based on Raman spectroscopy data that showed that the bulk water at a temperature of 310 K (37°C) contains water molecules with 3 and 4 hydrogen bonds in a ratio of 50/50%. Therefore, water molecules of the proteins’ first hydration layer having >2 hydrogen bonds do not change their configurational entropy upon transfer from protein surface to bulk water. The LEW content was calculated as the ratio of the number of LEW molecules to the total number of water molecules in the first hydration layer of a given area of the protein surface.

### ROC analysis

To obtain the optimal quantitative criterion for identifying the active sites of target proteins, ROC analysis using the maximum Youden index was used^19^,. Calculations were compared for known active sites, i.e., target protein surface regions that reliably bind ligands, and open surface regions, i.e., surface regions that reliably do not bind ligands. Inactive cavities were excluded from consideration, since their ability to bind any ligands *in vivo* is not precisely known. This allows us to obtain a cutoff threshold based only on those protein surface regions whose ability to bind or, conversely, not bind ligands is reliably known, thereby ensuring maximum sensitivity of the method.

## Data availability

The raw data generated in this study are available in the Zenodo repository https://doi.org/10.5281/zenodo.21222279.

## Acknowledgements

We thank Centre of Data Processing, Petersburg Institute of Nuclear Physics, NRC “Kurchatov Institute” for providing supercomputer facilities used in molecular modeling and MD simulations of proteins from representative set of human enzymes.

## Author contributions

S.P. and M.P. conceived the study. V.K. prepared the representative set of human enzymes. S.P. performed all the calculations and prepared software scripts and figures.

All authors interpreted the results. S.P. and M.P. wrote the manuscript. All authors approved the manuscript before submission.

## Additional information

### Supplementary information

The online version contains supplementary material available at […] Correspondence and requests for materials should be addressed to Svetlana Panasenko and Michael Petukhov.

---

**Supplementary Fig. 1.**
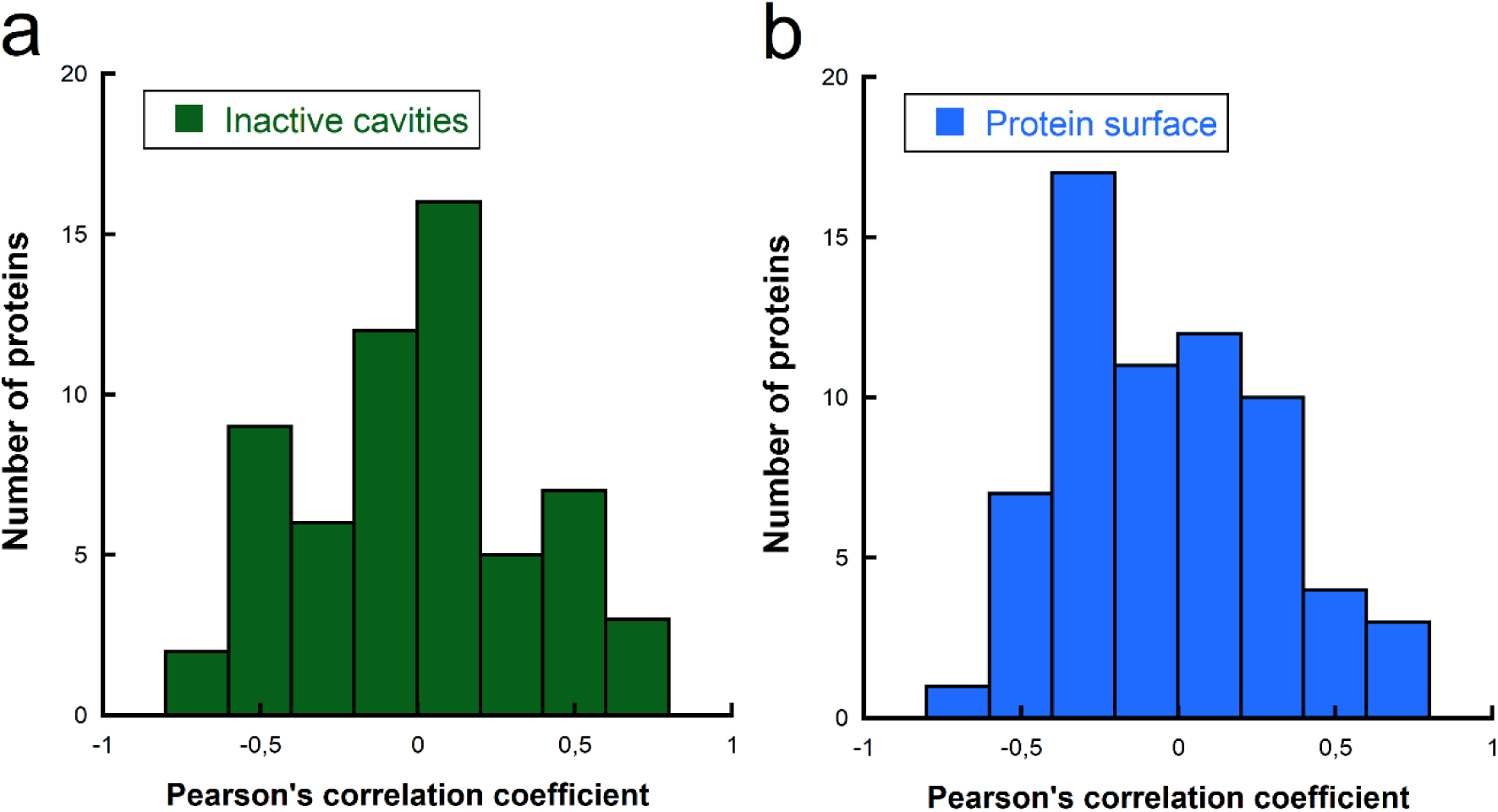
Distribution of Pearson correlation coefficients between the fraction of low-entropy water molecules and the fraction of hydrophobic ASA for control surface regions of the 65 proteins: **a**, inactive cavities; **b**, the rest of the exposed surface. The calculation was performed from ten equally spaced frames of the MD trajectory (analogous to Fig. 4b in the main text). The mean ± standard deviation values were: - 0.01 ± 0.37 for inactive cavities and - 0.04 ± 0.34 for the exposed surface.

